# Tracking Genomic Characteristics across Oceanic Provinces: Contrasting Early and Mature Plastic Biofilm Communities

**DOI:** 10.1101/2022.01.23.477364

**Authors:** Ryan P. Bos, Drishti Kaul, Erik R. Zettler, Jeffrey M. Hoffman, Christopher L. Dupont, Linda A. Amaral-Zettler, Tracy J. Mincer

**Affiliations:** Harbor Branch Oceanographic Institute, Florida Atlantic University, Fort Pierce, Florida, USA; Microbial and Environmental Genomics, J. Craig Venter Institute, La Jolla, CA, USA; Department of Marine Microbiology and Biogeochemistry, NIOZ Royal Netherlands Institute for Sea Research, Den Burg, Texel, The Netherlands; Department of Freshwater and Marine Ecology, Institute for Biodiversity and Ecosystem Dynamics, University of Amsterdam, Amsterdam, The Netherlands; Josephine Bay Paul Center for Comparative Molecular Biology and Evolution, Marine Biological Laboratory, Woods Hole, MA, USA; Department of Biology, Wilkes Honors College, Florida Atlantic University, Jupiter, Florida, USA

**Author notes:** Corresponding authors: Linda A. Amaral-Zettler, Tracy J. Mincer, **Email:**. **Author Contributions**: C. L. D., E. R. Z., L. A. A-Z., and T. J. M. conceptualized the research; E. R. Z. and J. M. H. collected samples; C. L. D., D. K., E. R. Z., L. A. A-Z., and R. P. B. were involved in data generation; C. L. D., D. K., E. R. Z., J. M. H., L. A. A-Z., and R. P. B. processed the samples; C. L. D., D. K., L. A. A-Z., and R.P.B. processed the data; R. P. B. and T. J. M. analyzed data; R. P. B. wrote the first draft of the manuscript. All authors edited and approved the final version of the manuscript. **Competing Interest Statement:** The authors declare no competing interests.

**Keywords:** bacteria, biofilm, early colonization, metagenomics, microplastics

## Abstract

While plastic has become omnipresent in the marine environment, knowledge of how plastic biofilm communities develop from functional metabolic and phylogenetic perspectives is nascent, although these data are central to understanding microbial ecology surrounding plastic substrates in the ocean. By incubating virgin microplastics during oceanic transects and comparing with naturally occurring plastic litter at the same locations, we constructed functional gene catalogs to contrast the metabolic differences between early and mature biofilm communities. Early colonization incubations were consistently dominated by Alteromonadaceae and harbored significantly higher proportions of genes associated with adhesion, biofilm formation, chemotaxis, defense, iron acquisition and utilization, and motility. Comparative genomic analyses with *Alteromonas, Marinobacter*, and *Marisediminitalea* metagenome assembled genomes (MAGs) spotlighted the importance of the mannose-sensitive hemagglutinin operon, adhesive genes genetically transposed from intestinal pathogens, for early colonization of hydrophobic plastic surfaces. Synteny alignments of the former operon also demonstrated apparent positive selection for *mshA* alleles across all MAGs. Early colonizers varied little in terms of large-scale genomic characteristics, despite the presence of latitudinal, salinity, and temperature gradients. Mature plastic biofilms, composed of predominantly Rhodobacteraceae followed by Flavobacteriaceae, that are critically important for carbon turnover in oceanic ecosystems, displayed significantly higher proportions of genes involved in oxidative phosphorylation, phosphonate metabolism, photosynthesis, secondary metabolism, and Type IV secretion. Our metagenomic analyses provide insight into early biofilm formation on virgin surfaces in the marine environment, as well as how early colonizers self-assemble, compared to mature, taxonomically, and metabolically diverse biofilms.

**Significance Statement:** Little is known about plastic biofilm assemblage dynamics and successional changes over time. Our results demonstrate that highly reproducible and predictable types of bacteria, with similar genomic characteristics, can initially colonize plastic in the marine environment across varying environmental gradients. The key gene sets involved in foundational bacterial colonization may have broad impacts for biofilm formation on plastic surfaces used in agriculture, biomedicine, environmental science, and food science. Genomic characteristics of early colonizers may metabolically underpin the origin of the ordered succession observed in marine microbial communities and be useful for predicting microbial community membership and biogeochemical processes.

## Introduction

Plastic marine debris is now recognized as an urgent threat to global ecosystems and has become ubiquitous in the ocean. Approximately 5.25 trillion plastic fragments weighing over 250,000 tons are afloat on the ocean’s surface (1), with microplastics (<5 mm) accounting for nearly 93% of the so-called ‘global particle count’. Since 2010, estimates indicate that between 4.8 to 12.7 million metric tons, or approximately 5% of annual plastic production is input to the global ocean each year (2) but could be as high as 11% of annual production, or 23 million metric tons (3). By the year 2050, the projected global plastic production is expected to surge to 1,500 million tons, nearly a four-fold increase from the current standard today (4), with limited infrastructure and policies in place to manage plastic waste.

Approximately 40-80% of archaeal and bacterial cells on Earth reside in biofilms, and these substrate-associated biofilms provide the foundation for many biogeochemical processes (5). Microplastic pollution presents a unique ecological niche in seawater, with the microbial community colonizing the surface of plastic, termed the ‘Plastisphere’ (6). Conservative estimates suggest that between 2.1 × 10^21^ to 3.4 × 10^21^ cells, or 1% of cells in the neuston layer alone colonize plastic globally and in biogeochemical terms, between 1,496 to 11,416 tons of carbon biomass reside on marine plastics, although these values are likely higher when including all ocean depths (7). Many plastisphere studies have amplified hypervariable regions of the 16S rRNA gene for archaeal/bacterial diversity, fewer studies have profiled exclusively eukaryotic or fungal diversity using the 18S rRNA gene locus or the ITS2 marker, and only five studies have used shotgun metagenomics from plastic to determine the functional composition of the Plastisphere (Table S1). Given the pervasiveness of microplastics in the marine environment, more functional metabolic data are necessary to gain a deeper understanding of the ecological implications of plastic substrates in the ocean. At the same time, plastic pollution stands as a global and evolutionarily recent perturbation that offers the prospect to study microbial ecological and evolutionary adaptations in near real time.

Regardless of sequencing methodologies, almost all previous plastisphere incubation studies have moored plastic particles to fixed locations and exposed these particles to local environmental conditions. However, the true nature of the Plastisphere is dynamic, spanning many environmental gradients, and therefore plastic represents a unique opportunity to study how metabolism and community membership varies on the same plastic particle across environmental gradients. Previous community relative abundance data (8-12) suggest that seasonal and biogeographical drivers facilitate the structuring of plastisphere communities, whereas other studies (12-14) suggest salinity and temperature as primary influences on microbial community structure. Importantly, the metabolic constraints for initial plastic colonization, as well as how these early colonizers impact secondary settlement on plastic remain an open-ended question.

Prior metagenomic studies have hinted that there are advantageous gene sets for a plastic-associated lifestyle (15, 16, 17), and few data exist on the beneficial gene sets used for colonizing the surface that plastic provides. While other studies have sampled DNA during the first week of colonization and from multiple time points to study community succession (12, 13, 18-23), only small-subunit ribosomal RNA gene sequencing was used to provide community profiling information. Consequently, the functional metabolic potential of early colonization of the Plastisphere remains poorly understood, although this information is critical for understanding general biofilm formation on hydrophobic surfaces. Though in different settings, the key themes of early colonization and biofilm formation on marine plastic surfaces may be applicable on plastics and other inanimate materials used in agriculture, biomedicine, and food processing facilities, which have been shown to harbor human pathogens (24).

In this study, we hypothesized that early colonizers of plastic in seawater, irrespective of surface sampling location across different oceanic provinces, would share large-scale genomic characteristics. To that end, we incubated virgin plastic particles along transects that were harvested incrementally underway to mimic an epiplastic community as it drifted through surface marine waters. These incubations were carried out alongside collections of free-drifting plastic particles, that served as proxies for mature biofilms from the same sampling locations. Using whole-genome-shotgun sequencing and a suite of bioinformatic tools, we established taxonomic and functional gene inventories for early and mature epiplastic biofilms. We further performed comparative genomic analyses with MAGs of the dominant early plastic colonizers to compare their genomic characteristics to describe the potentially advantageous gene sets for initial colonization and lifestyle on plastic.

## Results

Whole-genome-shotgun sequencing followed by preprocessing yielded ∼19,000,000 high-quality, paired-end reads per read library. Assembly information for all metagenomes can be viewed in Tables S2-S3. The supplement includes the heatmap containing the top 50 bacterial species (Figure S2) and taxonomic breakdown at the family level generated via Metagenomic Intra-species Diversity Analysis System (MIDAS; Figure S3) for aquarium seawater, cardboard, plastic, and wood communities. Figure 3 displays the Kaiju annotated relative abundances of bacterial and eukaryotic families observed for each respective sequence library from aquarium seawater cardboard, plastic, and wood. Aquarium seawater communities were replete with bacteria from the families Pelagibacteraceae, Prochloraceae, and Synechococcaceae, but also included members from Flavobacteriaceae and Rhodobacteraceae (Figure 3). Early epiplastic communities were dominated by Alteromonadaceae, which was composed of *Alteromonas, Marinobacter*, and *Marisediminitalea*, and in most of these read libraries, this family constituted >50% relative abundance on polyethylene (PE), polyhydroxyalkanoate (PHA), and polystyrene (PS). Given that Kaiju and MIDAS employ fundamentally different approaches (taxonomy based on sequence homology vs. curated phylogenomic trees), they provided a complementary and corroborative confirmation that Alteromonadaceae were the dominant early colonizers of plastic in this study. Flavobacteriaceae and Oceanospirillaceae were also present in most early colonization incubations. Thirty seven high-quality MAGs, including 14 *Alteromonas*, four *Marinobacter*, and eight *Marisediminitalea* MAGs were recovered from early colonization incubations, and have been annotated and described in Dataset S4. Multiple strains of *A. macleodii* (AD006, AD037, Balearic Sea AD45, Black Sea 11, English Channel 673) were detected on PE, PHA, and PS during the drifter experiment, yet *A. macleodii* strain Black Sea 11, which was highly abundant on PHA was rarely observed on PE and PS (Figure S2).

**Figure 1.**
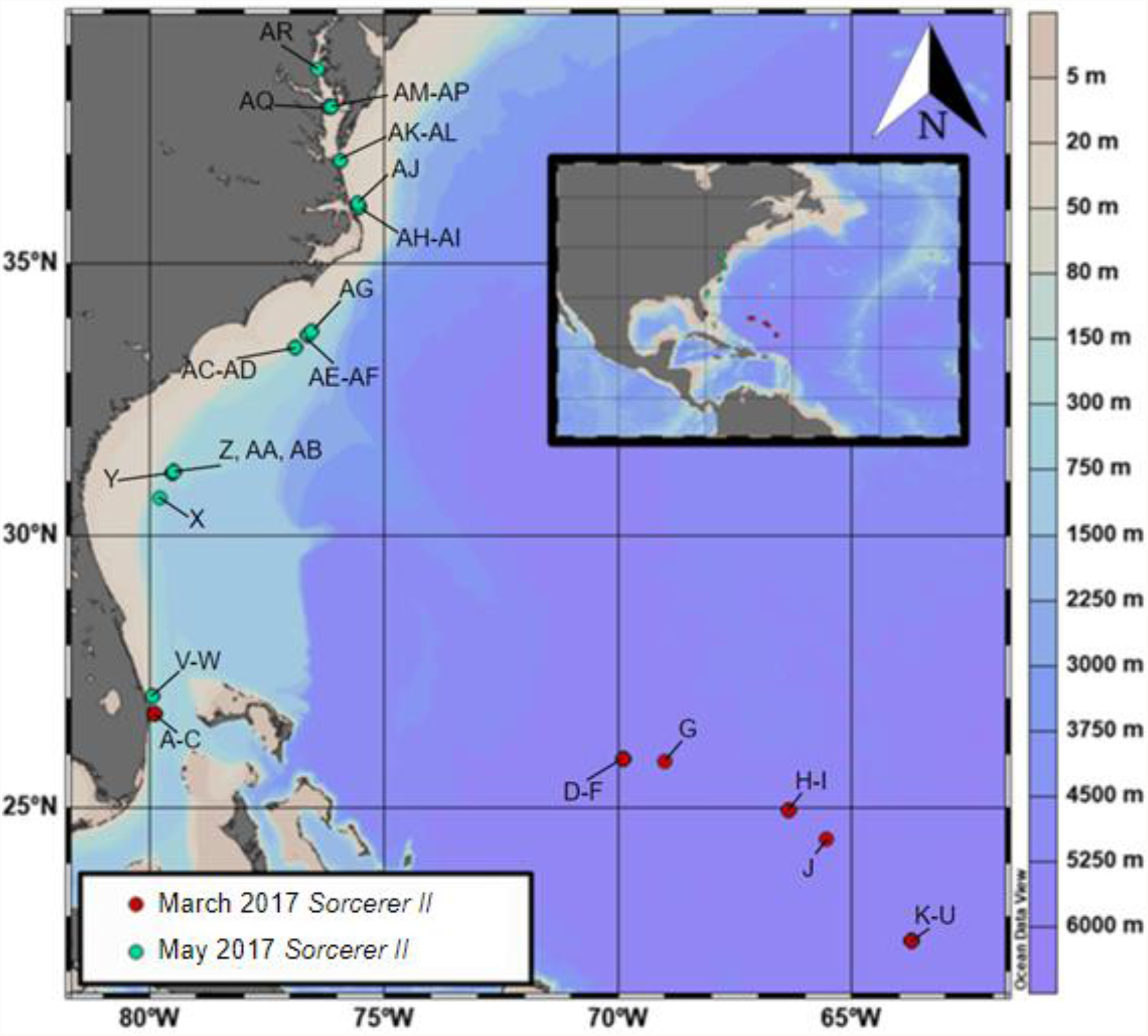
Map of sampling stations during the March 2017 (red shaded circles) and May 2017 (teal shaded circles) cruises aboard the *Sorcerer II*. Refer to Table S3 for corresponding sample labels.

**Figure 2.**
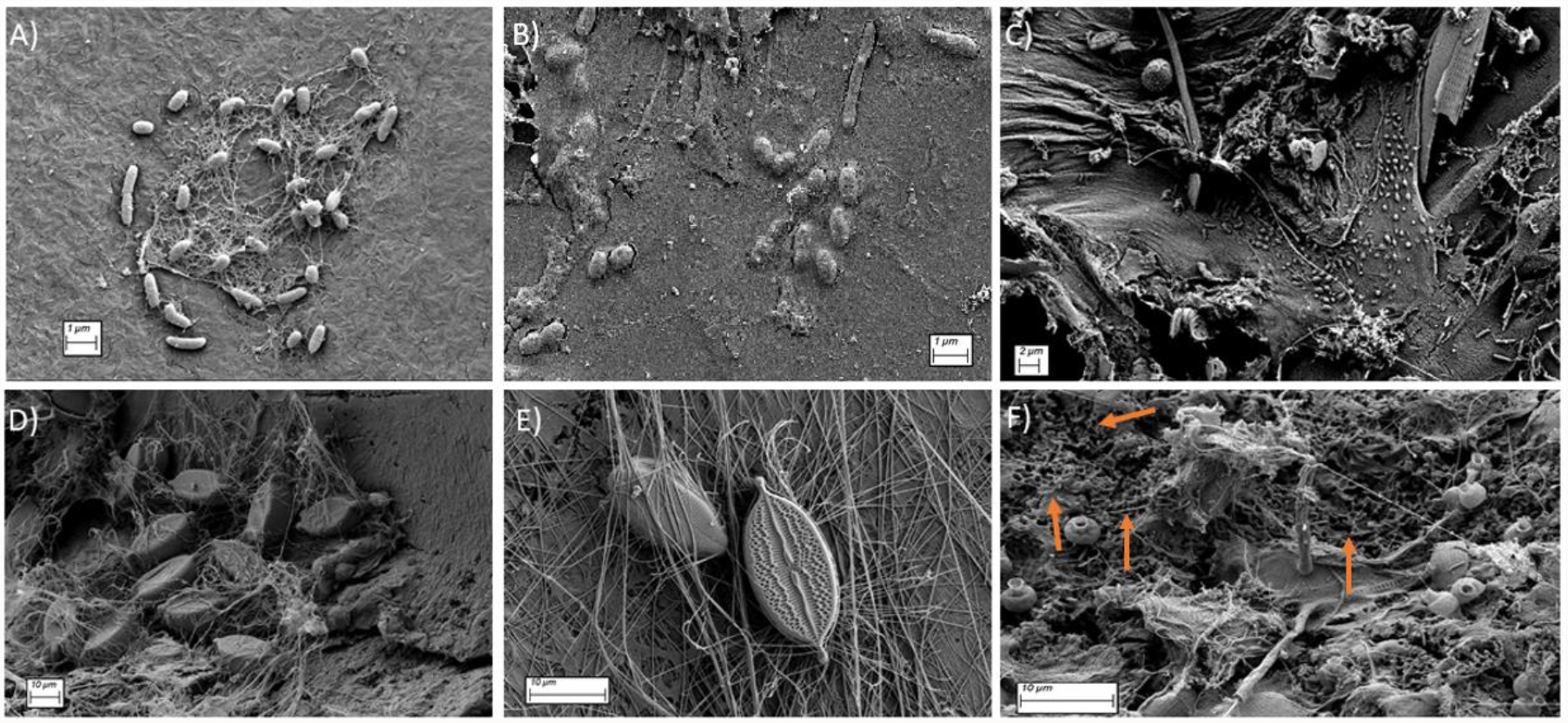
SEM images of early (A, B, C) and mature epiplastic biofilms (D, E, F): A) Microcolony of bacterial cells observed on PE. Lateral flagella and secreted pili are present, indicating irreversible attachment; B) various bacterial morphotypes embedded in the exopolysaccharide matrix on PS; C) early diverse biofilm on PHA consisting of different morphotypes of bacteria and eukaryotes; D-E) a cluster of diatoms entangled in secreted chitin ‘fingers’ that are used for attachment; F) the surface of highly weathered plastic collected via Manta trawl. Orange arrows denote areas where honeycomb-like, presumptive degradative pits are present.

**Figure 3.**
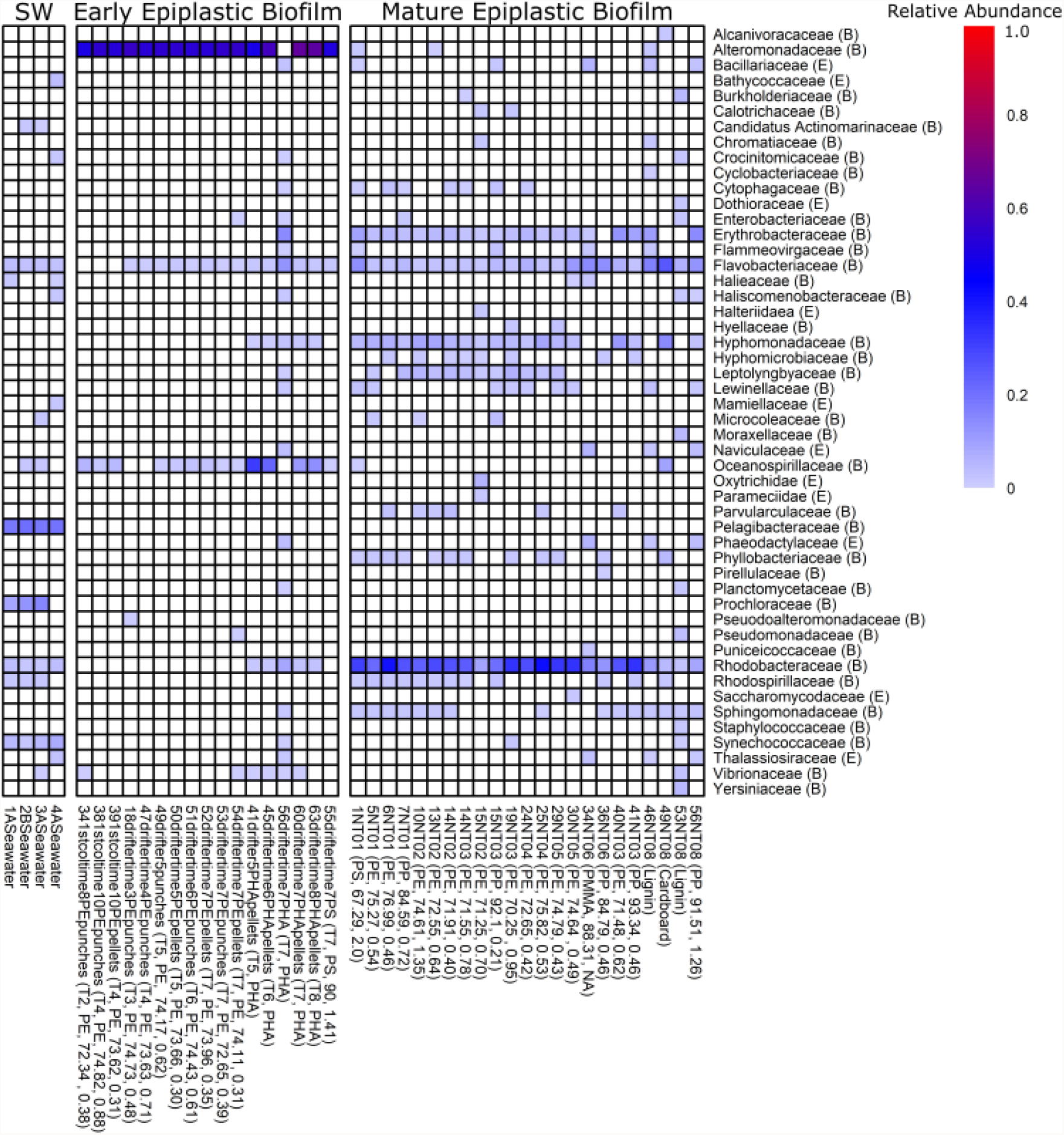
Taxonomic heatmap of bacteria (B) and eukaryotic (E) families annotated from aquarium seawater, cardboard, early and mature epiplastic, and wood communities using Kaiju annotations. SW indicates aquarium seawater metagenomic read libraries. Each column represents a unique read library. Relative abundances range from 0-1. PE, PHA, PS, PMMA, and PP indicate polyethylene, polyhydroxyalkanoate, polystyrene, poly(methyl methacrylate), and polypropylene, respectively. Read library names are displayed on the bottom portion of this figure followed by polymer type, percent match similarity with reference spectra, and carbonyl index.

Alteromonadaceae was rarely observed in aquarium seawater or mature epiplastic biofilm communities. Flavobacteriaceae and Rhodobacteraceae, which were present in all mature epiplastic biofilms, as well as Erythrobacteraceae and Hyphomonadaceae, were most abundant in these mature biofilms (Figure 3). There was a significant difference (PERMANOVA, P = 0.001) between the taxonomic composition of aquarium seawater and early and mature epiplastic biofilm communities (Figure S4), with all levels being statistically different from one another (post-hoc multiple comparisons, p-adj = 0.003, aquarium seawater vs. early colonization, 0.006, aquarium seawater vs. mature epiplastic biofilm, 0.003 early colonization vs. mature epiplastic biofilm). ANOSIM was used to discern the taxonomic resemblance between aquarium seawater and early and mature epiplastic biofilm communities and within each group, and these communities were significantly different from one another, but varied little within each respective group (ANOSIM, R = 0.9286, P = 0.001). Cardboard and wood samples were used for ordination but excluded from multivariate statistics because of the low sample size.

When grouping early and mature epiplastic communities and comparing with aquarium seawater communities, we observed significantly higher proportions of genes involved in the Clusters of Orthologous Groups (COG) categories Cell Motility, Defense Mechanisms, Function Unknown, General Function Prediction Only, Inorganic Ion Transport and Metabolism, Intracellular Trafficking, Mobilome, Signal Transduction Mechanisms, and Transcription in the epiplastic communities (Figure 4A). Eight categories were significantly more abundant in aquarium seawater communities, and these categories included Amino Acid Transport and Metabolism, Coenzyme Transport and Metabolism, Energy Production and Conversion, Lipid Transport and Metabolism, Nucleotide Transport and Metabolism, Posttranslational Modification, Replication, Recombination, and Repair, and Translation (Figure 4A). When disaggregating early colonizers and mature epiplastic biofilm samples, early colonization had a significantly higher proportion of genes associated with Cell Cycle Control, Cell Motility, Defense Mechanisms, Inorganic Ion Transport and Metabolism, Membrane Biogenesis, Signal Transduction Mechanisms, and Transcription (Figure 4B). In contrast, mature epiplastic biofilms had significantly more genes involved in Amino Acid Transport and Metabolism, Carbohydrate Metabolism, Coenzyme Transport and Metabolism, Energy Production and Conversion, Nucleotide Metabolism, Posttranslational Modification and Protein Turnover, Secondary Metabolite Biosynthesis and Metabolism, and Translation (Figure 4B).

**Figure 4.**
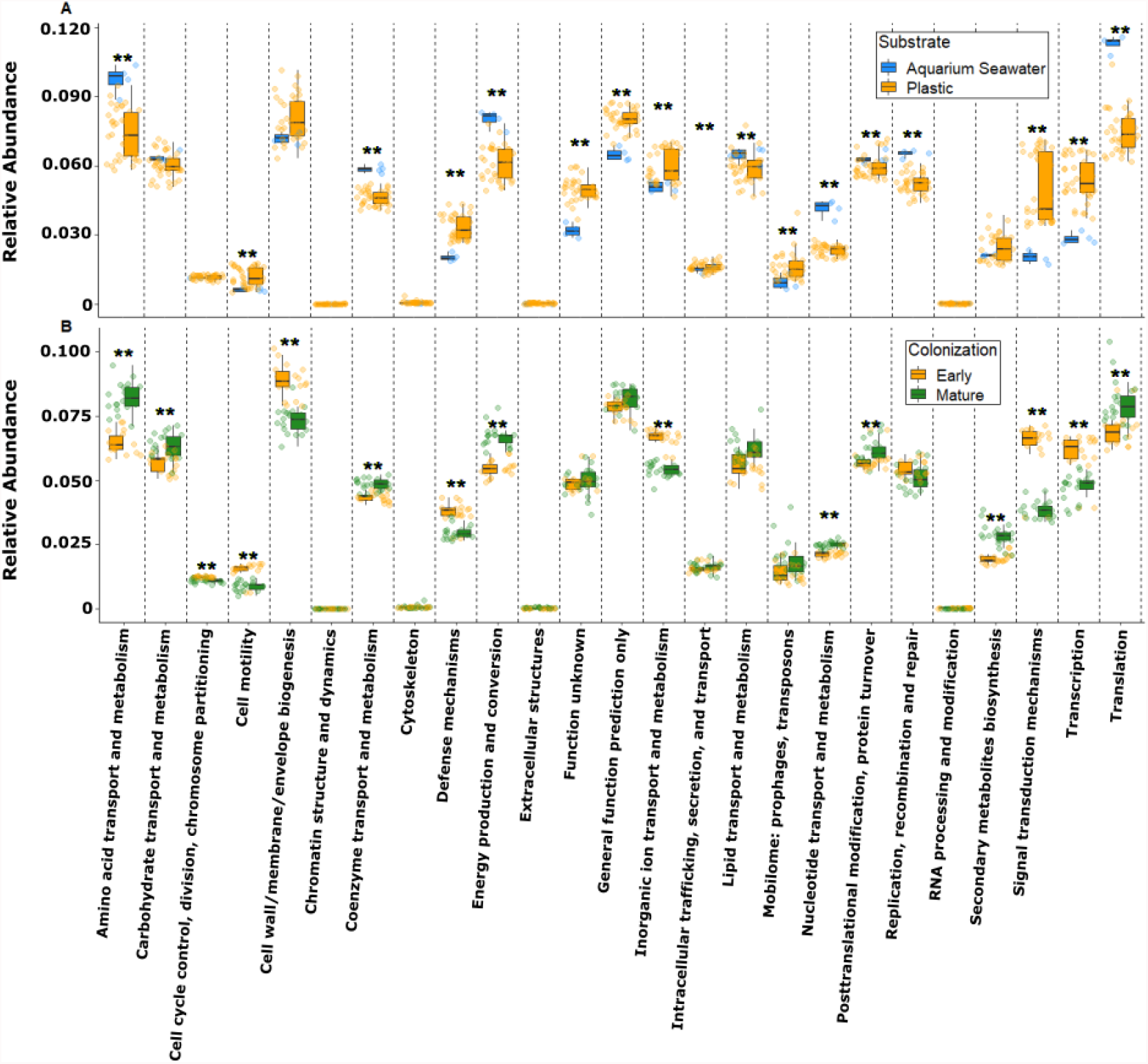
Comparison of COG categories by (A) substrate (aquarium seawater vs. plastic) and 1. colonization phase (early vs. mature) that were observed in aquarium seawater and epiplastic communities. ** Denotes statistical significance between substrates. Boxes represent the interquartile range, individual shaded circles represent one measurement for a category from one metagenome, solid horizontal black line is the median, whiskers are the maximum and minimum values.

When comparing functional metabolic potential between early and mature epiplastic biofilms and free-living communities there was a significant difference, with all communities being functionally distinct from one another (Figure S5A, PERMANOVA, P = 0.001, p-adj = 0.002 aquarium seawater vs. early colonization, 0.004 aquarium seawater vs. mature epiplastic biofilm, 0.002 early colonization vs. mature epiplastic biofilm). ANOSIM was used to discern the functional metabolic resemblance between aquarium seawater and early and mature epiplastic biofilm communities and within each group; these communities varied little within each respective group (ANOSIM, R = 0.7783, P = 0.001). When exploring functional potential observed in early and mature epiplastic biofilms, a total of 8314 unique KOfams were annotated, with 1452 and 1029 KOfams being enriched in early colonization and mature epiplastic biofilm communities, respectively (Figure S5B; Datasets S5-S6). When compared with mature epiplastic biofilms, early colonizers possessed enriched metabolic and regulatory characteristics involved with adhesion, biofilm formation, chemotaxis, environmental stress, iron utilization, motility, and pathogenesis (Figure 5). By contrast, in mature epiplastic biofilm communities, there was a significantly higher proportion of genes participating in oxidative phosphorylation, phosphonate metabolism, photosynthesis, secondary metabolism, and type IV secretion system pathways (Figure 6).

**Figure 5.**
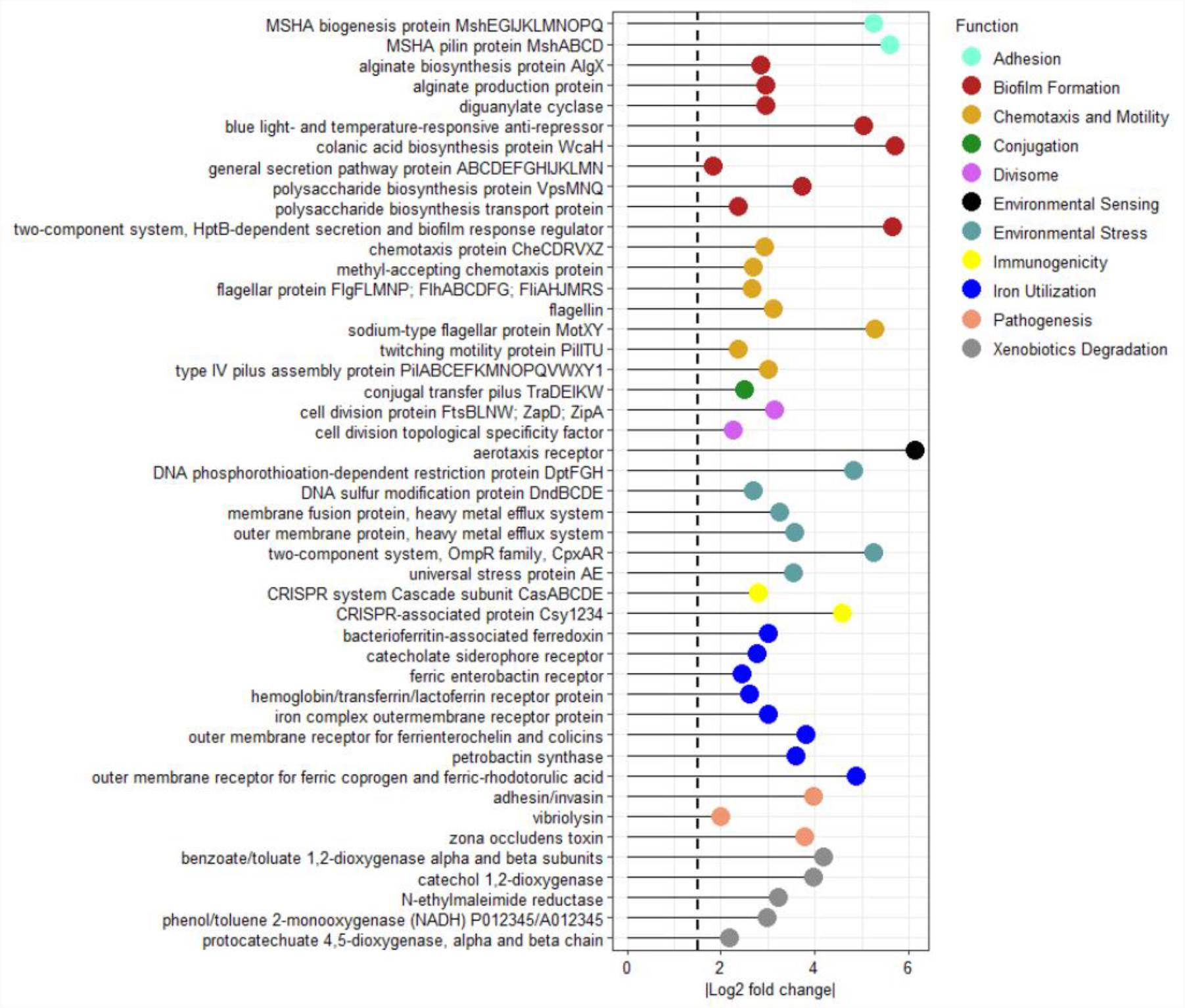
The mean Log_2_ fold changes of enriched KOfams with adjusted p-values (<0.05) and Log2 fold changes greater than the absolute value of 1.5 for early colonizers of plastic.

**Figure 6.**
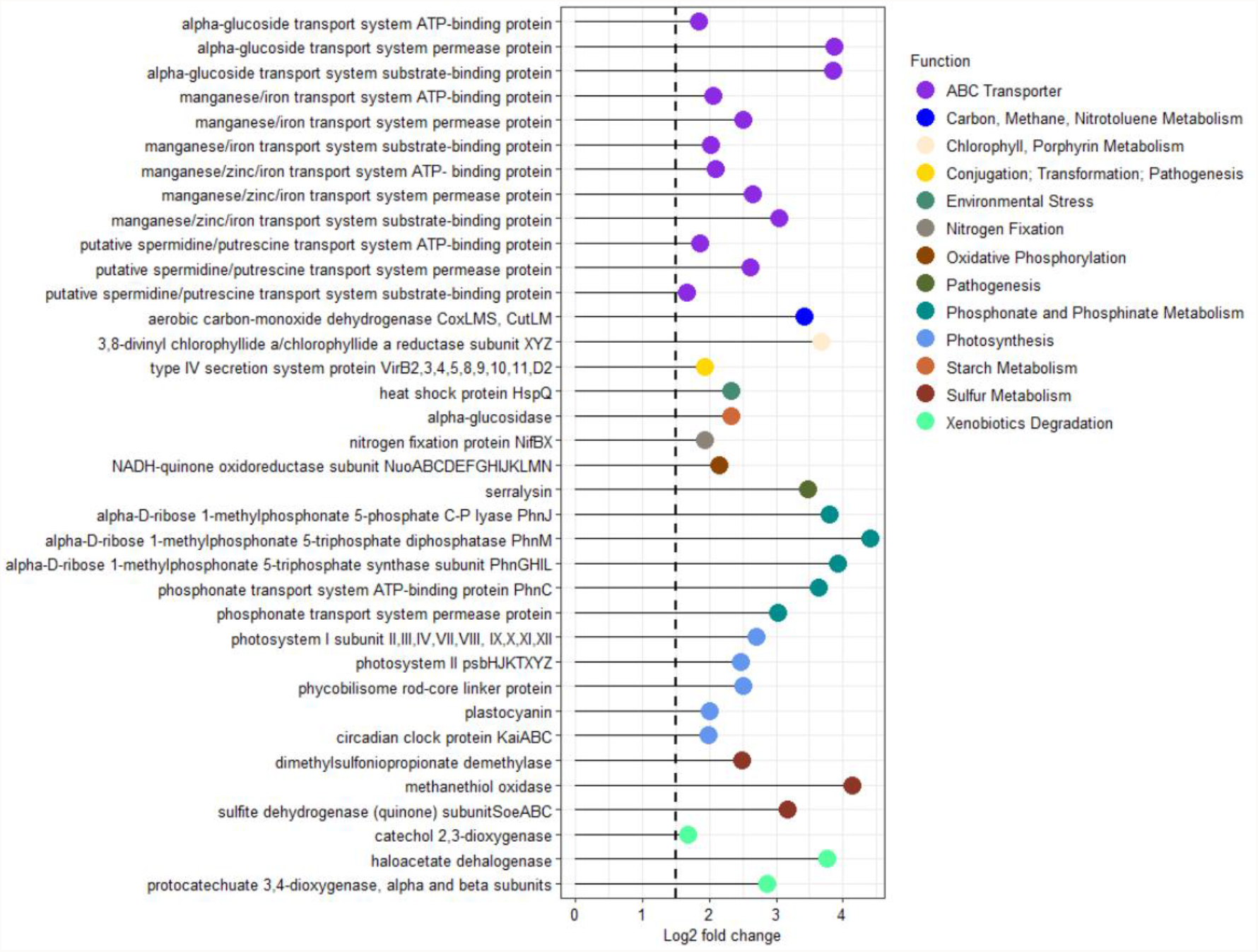
The mean Log_2_ fold changes of enriched KOfams with adjusted p-values (<0.05) and Log2 fold changes greater than the absolute value of 1.5 for mature epiplastic biofilms.

Using gene catalogs and read mapping, comparative genomic analyses with metagenomes and MAGs revealed that putative operons for the type IV mannose-sensitive hemagglutinin (MSHA) pilus were significantly enriched in early colonization and positively correlated with substrate hydrophobicity (Figure 5 and Datasets S4, S5, and S7). The same was true for *mshA*, except this gene was highly positively correlated with substrate hydrophobicity (Spearman’s correlation, ρ = 0.7573). Our *Alteromonas macleodii* MAGs recovered from PHA, which annotated as *Alteromonas* Kul 49, did not possess any components of the MSHA operon, whereas the *Alteromonas, Marinobacter*, and *Marisediminitalea* strains from PE and PS possessed full or fragmented portions of the operon, with some MAGs possessing multiple copies of *mshA*, each copy with a unique amino acid sequence (Datasets S4 and S7). *Alteromonas macleodii, Marinobacter flavimaris*, and *Marisediminitalea aggregata* synteny alignments of the MSHA operon and neighboring genes, that displayed high synteny and apparent positive selection for *mshA* can be viewed in Figures S6-S8.

## Discussion

The metagenomic data from plastics collected via Manta trawl and week-long shipboard PE, PHA, and PS incubations present a stark contrast between early and mature epiplastic biofilm communities (Figures 3-6). While we do not know the origin or age of environmentally-sourced plastics, carbonyl indices, SEM imaging, logistical logic, and our functional and taxonomic data suggest that environmentally-sourced plastics had older biofilms. Potential caveats to this generalization include perturbations to a biofilm community such as a surface grazing, ingestion by organisms and subsequent egestion, as well as physical abrasion that may obfuscate the exact age of a biofilm community. Alteromonadaceae dominated early colonization of the plastisphere in 16 of 17 PE, PHA, and PS drifter read libraries (Figure 3), but none of the mature biofilms on net captured particles, and in the present study, this family was composed of genera *Alteromonas, Marinobacter*, and *Marisediminitalea. Alteromonas* spp. are cosmopolitan and highly transcriptionally active members of particle-associated communities in the marine environment (25, 26), also observed on ship-hulls (27), transparent exopolymer particles (28), and marine snows (29). The presence of Alteromonadaceae has been noted in early epiplastic biofilms (12, 13, 19, 30), with Alteromonadaceae being used to discriminate colonization between aliphatic and aromatic polymers (31). However, comparing the extent of colonization here with other 16S rRNA gene amplicon-based early colonization studies (12, 13, 18-23) should be done with caution because of the fundamental differences between amplicon and whole-genome-shotgun sequencing methodologies. Experimentally determining that Alteromonadaceae members are consistently the major early colonizer across polymer types, with high agreement between Kaiju (taxonomy based on sequence homology) and MIDAS (taxonomy based on curated phylogenomic trees) is notable. The Alteromonadaceae genera *Alteromonas, Marinobacter*, and *Marisediminitalea* have large flexible, genomes, and are perhaps better suited to colonize diverse substrates and metabolize eclectic carbon sources due to harboring a broad metabolic repertoire (26, 32). These three genera also include hydrocarbonoclastic copiotrophs (33, 34), attracted to dissolved organic matter (35), and their dominance on plastic may be due to (a) rapid chemotaxis towards additives, hydrocarbon leaching (36, 37), or nutrient pulses emanating from plastic polymers; (b) accumulation of hydrolysable macromolecules on plastic’s surface from the so-called Zobell Effect (38); (c) cooperative metabolic incentives, defense, and increased growth and reproduction from colonizing a substrate. Targeted microfluidic assays with plastic dissolved organic matter are still required to verify whether leached additives and hydrocarbons lure microorganisms.

We hypothesize that metabolically predictable types of hydrocarbonoclastic bacteria that harbor a large, versatile genome initially colonize plastic in the marine environment. Based on high consensus between our COG20 categories and functions, KOfam, and Pfam annotations, we posit that early colonizers of marine surfaces possess similar large-scale genomic characteristics such as the flagella and pili involved in accelerated adhesion, biofilm formation, chemotaxis, and motility (Datasets S4 and S7). Functional gene inventories are missing from prior small-subunit ribosomal RNA gene sequencing samples of early plastic colonization, but the genome characteristics of primary early colonizers from those studies appear to support our hypothesis. For example, the presence of *Aestuariibacter*, an *Alteromonas*-like microorganism, *Oleiphilus*, and a *Roseobacter*-like bacterium on PE after two days of incubation were reported (22). *Oleiphilus* spp. specialize in biodegradation of aliphatic hydrocarbons and appear to have more of their proteome allocated to cell motility and environmental sensing when compared with early colonizers in the present study, and *O. messinensis*, for example, has a genome larger than 6 Mb (39). The *Roseobacter* group has genomes that average 4.4 Mb (40), and this clade has members that metabolize aromatic compounds (41), including lignin (42), which may be useful for breaking refractory ether bonds in phenolic plastics (42). Gammaproteobacteria, including Alteromonadaceae, followed by the Alphaproteobacterial family Rhodobacteraceae, predominantly *Roseobacter*, were abundant on PE and PS after seven days of incubation (12). Like Alteromondaceae, Rhodobactereaceae is frequently reported to be an early colonizer of biotic (43) and abiotic (44, 45) marine surfaces, as it is metabolically capable of rapid responses to resources (46) and has been purported to be critically important for facilitating initial biofilm formation and colonization by other bacterial species (47). Rhodobacteraceae has also been implicated in oil remediation post Deepwater Horizon (35). Oceanospirillaceae (20-30%) and Alteromonadaceae (14%) dominated in terms of relative abundance on plastic after one day of immersion, with Rhodobacteraceae (21-44%) becoming most relatively abundant after four days (21). Oceanospirillaceae, which was composed mainly of the large genome *Oleibacter* (∼3.8 Mb), is hydrocarbonoclastic (48) and possesses giant flagellins that enhance the thickness of the flagella (49). These observations suggest that *Oleibacter* is heavily invested in motility, with the increased flagellar thickness potentially being a mechanical adaptation to allow quicker or more efficient swimming (49). Saprospiraceae followed by Flavobacteriaceae were most abundant on six different polymer types after one week of incubations in the Caribbean (20). Saprospiraceae are noted to produce copious exopolysaccharides (50) and have been observed in activated sludge (51), with these microorganisms being likely to degrade complex carbon sources (52). Flavobacteriaceae are also known to derive energy from diverse organic materials (53), with *Flavobacter* being a known degrader of Nylon-6 (54). Flavobacteriaceae possess a gene set for rapid gliding motility (55), which could explain their potentially important role for carbon turnover in epiplastic communities. Both Flavobacteriaceae and Saprospiraceae have been implicated for oil remediation following the Deepwater Horizon oil spill (56). Observations from the present study and all early colonizer incubations are important because it appears that the bacteria that initially colonize plastic are phylogenetically diverse yet share common genomic traits. These shared traits may be useful to understand the type of community that initially colonizes plastic and the community that succeeds it.

For years, it has been hypothesized that there is a forecastable community succession on virgin surfaces in the marine environment, with bacteria serving as pioneering settlers, followed by microalgae such as diatoms, and invertebrates as the biofilm matures (38, 57, 58). Our first samples were taken after 2 days of colonization, so we cannot say that bacteria are the first colonizers, but 16 of our 17 early colonization read libraries had bacterial reads accounting for >99% of mapped reads across all time points (Figure 3), so they dominated our early colonization samples. While tempting, our data cannot be used to determine whether a stable ‘climax’ community is ever reached, or if functional community metabolism converges over time regardless of the surface and physicochemical properties of different polymer types, although our mature biofilm communities separated by substrate may provide some insight. In our mature biofilm communities, there was no significant difference in the functional KOfam matrix between polymer types and substrates (ANOSIM, P = 0.002, R = 0.4493; PERMANOVA, P = 0.132, R2 = 0.32041), which suggests, albeit with limited sample size, that mature biofilm communities may functionally converge (Figure S9). Interestingly, for day 7 PHA in March, diatom families, which included Bacillariaceae, Phaeodactylaceae, and Thalassiosiraceae accounted for 8% of annotated reads. These observations suggest that a different type of community, which taxonomically and functionally resembled a mature epiplastic biofilm was present on the day 7 PHA samples during the March cruise and support the hypothesis that some epiplastic communities could become mature in as little as nine days (note single orange dot representing PHA day 7 sample interspersed with green mature biofilm samples, Figures S4B and S5A; 59). Datta et al. (60) present an alternative scenario that could also explain the presumed rapid succession changes from early biofilm to mature epiplastic community. In that study, the authors demonstrated that chitin-attached communities undergo rapid successions, whereby motile, particle-degrading taxa are selected for during early successional stages (24-48 hours). The authors further argue that when secondary consumers that cannot use the substrate as a carbon source colonize, community metabolism shifts away from the substrate. It is not immediately clear whether Datta’s 2016 observations on chitin can be applied to a synthetic, recalcitrant polymer like PE, or PS, but, perhaps, it can be applied to a biodegradable plastic like PHA.

Metagenomic and metaproteomic studies to date suggest it unlikely that microorganisms are holistically degrading fossil-fuel based plastic in the ocean due to the presence of labile compounds in epiplastic biofilms that could be prioritized over recalcitrant plastic carbon (17). However, several genes putatively involved in hydrocarbon degradation observed in the present study insinuate that microorganisms may play some role in degrading plastic in the marine environment (Figures 5-6; Dataset S4), although the microorganisms responsible and the rate at which microbial degradation occurs remain elusive. For example, the presence of micron sized honeycomb-like pits on the plastic’s surface suggest that microorganisms are degrading plastic in the marine environment (Figure 2F). The presence of hydrocarbonoclastic bacteria initially colonizing plastic (particle-degrading specialists), and the shift to Flavobacteriaceae and Rhodobacteraceae in mature epiplastic communities, which are critically important for carbon turnover in the ocean (secondary consumers), hints that plastic communities may follow a similar multiphasic trajectory as chitin communities (60). The constraints, duration, and presence of these phases in epiplastic communities are not currently understood but may provide insight into the extent to which plastic is degraded, how much plastic carbon contributes to the recalcitrant carbon pool in the ocean (61), as well as the trajectory of the community.

Enriched COG categories during early colonization were complemented by KOfam annotated genes for adhesion, biofilm formation, chemotaxis, environmental sensing and stress tolerance, hydrocarbon degradation, iron utilization, and motility (Figures 4-5). Methyl-accepting chemotaxis proteins, which are transmembrane receptors that induce a chemotactic response, were accompanied by the Che operon, which acts as a flagellar switch to direct movement towards stimuli (62, 63). Genes involved in flagellar assembly were significantly enriched in early colonization, and these genes have been noted as important for nutrient acquisition (64). While motility is a metabolically expensive trait, it has been shown to confer a selective advantage to microorganisms who invest in genes involved in these abilities (64). This observation paired with the significantly higher proportion of aerotaxis receptors in early colonization suggests that these bacteria have a competitive advantage for nutrient acquisition. Bacterial motility is also reported to provide an advantage when in the presence of bacteriophage (65). The significantly higher abundance of the COG categories ‘Cell Motility’ and ‘Defense Mechanisms’, KOfam annotated CRISPR-associated complexes, and phage shock proteins in early colonization suggests that early colonizers may be the target of bacteriophage (Figures 4-5 and Datasets S4-S6, 66), perhaps in their planktonic life stages. Few data exist on how viral interactions shape the plastisphere and should remain a topic of future research.

The MSHA biogenesis and pilin proteins are critical for structural assembly, surface adhesion and selection, and are best characterized in pathogenesis by *Vibrio cholerae* (67). These genes also appear to play a role in attachment to eukaryotes (68) and zooplankton (69). The MSHA operon was significantly enriched in early colonization and was positively correlated with substrate hydrophobicity, or water contact angle (PE water contact angle, θ = 101.7°, 70), PS (θ = 87°, 71), and PHA (θ = 75°; 72), which underscores that surface properties of plastic polymers including hydrophobicity, surface roughness and chemical structure are potential barriers for early colonization (Figure 5, 13, 73). Hydrophobicity has also been correlated with cellular abundance on plastic surfaces after week-long incubations (8), but no correlation was observed after eight weeks, and this suggests that physicochemical properties have limited influence after initial colonization, and that general biofilm processes and microbial interaction processes instead of polymer-specific properties shape mature epiplastic communities.

The observation that *Alteromonas macleodii* strain Black Sea 11 was rarely present on PE and PS samples but highly abundant on PHA, further underscores the importance of MSHA for early colonization of highly hydrophobic surfaces. Strain Black Sea 11 does not possess MSHA, and the same is true for the *A. macleodii* MAGs binned from PHA, which annotated as *Alteromonas* Kul 49, a species that also does not possess the MSHA operon (Figure S2 and Datasets S4 and S7). Importantly, a MAG does not represent clonal individuals, but rather a group of closely related lineages, in this case *Alteromonas* spp., that gives insight into that genus’ metabolism. The observation that MSHA was not observed in any *Alteromonas* MAGs binned from PHA is striking. These data suggest that possession of MSHA and multiple copies of *mshA* may provide a competitive advantage for early colonization of highly hydrophobic surfaces. What remains unclear, however, is whether microorganisms without MSHA can directly and competitively colonize hydrophobic plastic surfaces like PE and PS or are required to coordinate with earlier colonizers that do possess MSHA and colonize secondarily on top of secreted exopolysaccharides. Gene-expression and genetic manipulation studies with model systems are necessary for understanding MSHA’s role for early plastic colonization and will provide important implications for how microbial communities self-assemble on plastic.

While there appear to be predictable types of bacteria that initially colonize plastic in the marine environment, the possibility of a universal phenotype emerging for plastic colonization has not been explored. We argue that a broadscale and geologically young perturbation event such as ubiquitous plastic pollution presents an unprecedented opportunity to study evolutionary adaptations to novel surfaces in near real time. The MSHA virulence machinery is reported to be acquired as a mobile genetic element (74), with the genes directly flanking the operon, the rod-shape determining proteins *mreBCD* on one end and a single-stranded DNA binding protein on the opposite side being observable in some of our MAGs (Dataset S4). By leveraging synteny alignments of the MSHA operon between each clade’s closest reference genomes and our *Alteromonas macleodii, Marinobacter flavimaris*, and *Marisediminitalea aggregata* MAGs that were binned from plastic metagenomes, there appears to be high synteny, as well as apparent positive selection for alleles of *mshA* across all species that possessed the operon for reliable comparisons (Figures S6-S8). For microorganisms exhibiting a predilection towards plastic, it is plausible that genes involved in attachment may be positively selected for. Acquisition of the MSHA operon by open ocean bacteria may provide a competitive advantage for colonization of hydrophobic surfaces and may be a possible adaptation to the new microenvironment that plastic surfaces present. Of equal interest is the observation that some of our MAGs possess multiple copies of *mshA* that have fundamentally different amino acid sequences (Datasets S4 and S7). Some *mshA* gene products may be limited by chemical and physical constraints, towards certain amino acid sequences, for example in the MSHA binding domains, that are optimized for colonizing different surfaces, with each copy being predilected towards specific surfaces and interactions with microorganisms.

Diguanylate cyclase enzymatically triggers creation of cyclic-di-GMP, which is an intracellular signaling molecule that is critical for biofilm formation and persistence (75) and in our results this was supplemented with alginate and colanic acid biosynthesis genes, as well as several other genes for polysaccharide biosynthesis and general secretion of exopolysaccharides (Figure 5). Cyclic-di-GMP has also been shown to be a critical messenger for transitioning between motile and sessile stages (76). Interestingly, the blue light and temperature responsive anti repressor was enriched in early colonization, which implies that these microorganisms may secrete more exopolysaccharides when exposed to lower temperatures and higher levels of ultraviolet light (77). While not encountered on our cruises, *Alteromonas* spp. biomass has been reported to be absent in cold environments (<10°C, 78), and it is plausible to hypothesize that the epiplastic biofilm may enable cells to persist across environmental gradients in otherwise inhospitable environmental conditions. The same could be true for other abiotic factors like salinity. For example, Oceanospirillaceae, a family of halophilic microorganisms, displayed a presence suggesting potential activity, although their relative abundance was observed to decrease from 31% to 13.37% on PHA as it drifted across a decreasing salinity gradient in Chesapeake Bay (Figure 3; Table S3).

The significantly higher abundance of iron receptors and utilization genes in early colonization of the plastisphere is of particular interest because *Alteromonas* and *Marinobacter* produce siderophores (79), iron chelating organic molecules that serve to transport iron across the cell membrane (Figure 5). Reductive evolutionary theory states that a neighboring microorganism may experience vital gene loss through natural selection if other community members have leaky metabolic pathways that can be used as public goods, whereby microbial cheaters can still obtain quintessential nutrients (80). Siderophores provide a prime example of this phenomenon, also known as the Black Queen Hypothesis (81), as they can be exploited by community members if they still possess the compatible transporters (80). For example, the *A. macleodii* MAGs binned from PE and PS possessed the gene petrobactin synthase (82, 83), whereas the *A. macleodii* MAGs that were binned from PHA and found in low abundance on PE and PS did not have this gene present, and neither did their closest related type strain. All of the PE, PHA, and PS colonizing *A. macleodii* strains, as well as *Marisediminitalea* MAGs possessed catecholate siderophore and TonB receptors (Dataset S4), irrespective of siderophore producing ability or lack thereof. These observations, paired with significantly higher abundance of iron utilization genes involved in siderophore production and utilization during our drifter experiment suggest that early colonizers such as *Alteromonas* and *Marinobacter* may incentivize future colonization of plastic, as well as aid in overcoming iron limitation (common in oligotrophic surface waters), and therefore may be harbingers of the succeeding biofilm community.

In the present study, highly reproducible and phylogenetically distinct, hydrocarbonoclastic bacteria sampled across different oceanic provinces possessed similar large-scale genomic characteristics. While functional annotations are missing from previous 16S rRNA gene early plastic colonization studies, the traits of early colonizers observed in those studies also appear to support our hypothesis (12, 13, 18-23). Therefore, early colonizers appear to have larger, versatile genomes that are accentuated by either multiple copies of genes used for, or complete pathways involved in chemotaxis and environmental sensing, motility, attachment, and biofilm formation. A clear shift in metabolic potential was observed between early and mature plastic biofilms, which had significantly higher proportions of genes involved in oxidative phosphorylation, phosphonate metabolism, photosynthesis, secondary metabolism, and Type IV secretion (Figure 6), and critically, a significant reduction in the proportions of genes that appeared to be important for initial colonization. These data suggest that metabolic characteristics can be used to differentiate biofilm stages, and more data are necessary to analyze whether initial metabolic characteristics on substrates can be used to predict community succession. Based on the presence and high abundance of many *Alteromonas* strains observed in the same early colonizer communities (Figure S2), it is likely that there are coordination mechanisms and interactions and metabolic complementation between strains that are further driving community assembly and functioning. Future studies implementing metatranscriptomics would be useful as this could provide evidence of a) what microorganisms/genes are transcriptionally active in early and mature biofilms; b) which genes are differentially expressed between intrageneric strains; and c) how gene expression varies with time on a surface and with secondary settlement.

## Materials and Methods

### Sample Collection

The Drifter experiment was developed to simulate drifting plastic particles in surface marine waters and sample the associated epiplastic microbial community along a marine transect at different time points. Aquarium seawater and plastic samples were collected during two cruises aboard the R/V *Sorcerer II* from West Palm Beach, FL to Marigot on the island of St. Martin (March 19th-26th, 2017) and West Palm Beach, FL to Annapolis, MD (May 14th-18th, 2017; Figure 1). Samples were collected in two different ways: 1) shipboard incubations (2-7 days), which were incubated underway and harvested incrementally across the cruise track; and 2) Manta trawls (333 µm) that were deployed alongside locations where the short-term incubation samples were harvested. The short-term incubations represented the early colonization component of this study, whereas the particles collected via Manta trawl represented a proxy for mature biofilms collected at the same sample locations.

The short-term shipboard incubations consisted of incubating PE “Infant Water” jug punches from Walmart, pre-industrial PE pellets (Dow), PHA pellets (Meridian Holdings), and PS pellets (source unknown) in a 4-L precleaned aquarium aboard the ship, and regularly replenishing the aquarium with surface seawater along the transect for one week. The March and May Drifter experiment set ups included ∼200 circular ‘punches’ from the high-density PE jug, ∼200 PE pellets, ∼100 PHA pellets, and ∼100 PS pellets. All plastic particles and the aquarium were precleaned in 10% bleach then rinsed in distilled water for 30 minutes, followed by three rinses with laboratory grade deionized water. A 20-L plastic bucket was also precleaned and used as a reservoir for the experiment. For the March cruise, plastic particles were loose within the aquarium, and seawater in the 20-L bucket was pumped continually through the aquarium and replaced with surface seawater approximately every two hours during the day. For the May cruise, a slow flow-through seawater system was developed to continually add surface seawater to the 20-L bucket. The aquarium was covered with 1-mm fiberglass netting to retain the plastic in case of rough seas, and to provide some shading. Starting at two days, approximately 10 particles of PE, PHA, and PS punches and/or pellets were harvested using sterile forceps and preserved for DNA extraction and imaging (see below). Using a peristaltic pump, 2L of aquarium seawater from the 20-gallon incubator bucket was filtered through 0.22 µm Sterivex filters (Merck, Germany) periodically during the short-term incubations to compare the epiplastic community on the experimental pieces with cells in the surrounding aquarium seawater or that might be sloughing off plastic surfaces in the bucket.

Samples were further categorized based on the source (aquarium seawater, cardboard, plastic, and wood), as well as the colonization phase of microorganisms (free-living; early and mature biofilms). The measured carbonyl index of each plastic along with the appearance of each plastic’s surface (e.g., thick biofilm) were used to differentiate early from mature epiplastic biofilms. More detail is provided in section titled ‘Oxidation Degree’. Environmental data, including phosphate, nitrate, nitrite, silica, oxygen, salinity, and temperature were collected in conjunction with plastic sampling (Table S3).

### Sample Preservation

At each time point, incubated microplastics and plastics collected with Manta trawls were preserved for both molecular analysis and imaging. Multiple punches and pellets were pooled for each purpose to increase the amount of biofilm available, while net samples were cut into two pieces for molecular analyses and imaging. In each case the samples for molecular analysis were immediately transferred to Puregene cell lysis solution (Qiagen, Valencia, CA) and frozen at -20 °C for downstream whole-genome-shotgun sequencing. Samples for imaging were fixed in 4% paraformaldehyde (EMS, PA) then transferred to a phosphate buffer/ethanol mix and stored at - 20 °C till prepared for SEM imaging. SEM sample preparation and imaging followed the approach described in Zettler et al. (6), except a Leica EM CPD300 was used for critical point drying. Sterivex filters were flooded with Puregene cell lysis solution, sealed, and frozen at -20 °C.

### Polymer Identification

Polymer identification was achieved by using attenuated total reflectance in the spectral range from 3600-1250 cm-1 on an in-house Fourier Transform Infrared Spectrometer (Thermo Scientific™ Nicolet™ iN10, USA). While polymer identification on post-DNA extracted material is not ideal for quantifying oxidative state, all samples were treated equally and were therefore considered comparable. The polymer type of each particle was determined by comparing its chemical spectrum against the Hummel Polymer Library using OMNIC software (Thermo Fisher Scientific, USA). Auto baseline correction, atmospheric removal, and spectral optimization parameters were applied to each spectrum. Percent similarity, calculated by OMNIC Picta software interpretation was substantiated in one of two ways: 1) if there was greater than a 70% match with reference spectra; or 2) spectra with a match between 60% and 70% were reinterpreted by confirming the presence of specific diagnostic polymer peaks. The particle was determined to be plastic if the highest matching polymer type was observed during initial and secondary diagnostic steps.

### Oxidation Degree

The oxidation degree of plastic was measured concomitantly with polymer typing. Based on the application, plastics are manufactured with additives (e.g., plasticizers, scavengers, stabilizers) and consequently experience varied levels of chemical transformation upon exposure to environmental conditions such as temperature and ultraviolet light. The oxidation of plastic was calculated by measuring the ratio of the absorbance band area of the carbonyl group at 1630-1850 cm^-1^ and the olefinic band area at 1420-1490 cm^-1^ (84, 85). Carbonyl indices were not taken from PHA incubations due to the presence of a diagnostic peak found in the same region as the carbonyl band.

### Bioinformatic Pipeline

Total genomic DNA was extracted using the Gentra Puregene Tissue DNA Isolation kit (Qiagen, Valencia, CA). DNA extractions were performed using a modified bead beating approach described in (9). Library preparation was done using the Swift2S (Swift Biosciences INC., Ann Arbor, MI) library preparation kit following the manufacturer’s instructions, and DNA was sequenced using whole-genome-shotgun sequencing on a 150 base pair paired-end Illumina NextSeq at the J. Craig Venter Institute, La Jolla, CA. Raw reads were preprocessed to remove ligated adapters, regions of low complexity, and low-quality sequences using fastp (v0.12.5), with default parameters (86). PolyG tail trimming (--poly_g_min_len 50) and a quality threshold (--qualified_quality_phred 12) were used, and the quality of preprocessing steps was assessed using FastQC (87). The freely available software and data platform KBase (KBase, http://kbase.us/) was leveraged to assemble our quality-controlled, paired-end sequences using metaSPAdes (88), with kmers [21, 33, 55] and a minimum contig length of 500 nucleotides.

A Unix PowerShell script was created to take all metagenomic assembly files (.fa) in a working directory and enumerate gene counts for all genes in each assembly to construct a composite gene catalog. This script is available at https://github.com/Echiostoma. Briefly, each metagenomic assembly was iteratively converted into an Anvi’o (v7.0 dev, 89) compatible contig database using the anvi-gen-contig-database command. Open reading frames were predicted using Prodigal (v2.6.3, 90) and putative genes were annotated using COG20 Categories and Functions, KOfams, and Pfams with an e-value threshold of 1e^-4^ (filtered with awk), which is equivalent to a 0.001% chance that a gene is assigned to the wrong family. Annotated genes for each assembly were iteratively exported using the anvi-export-functions command. The exported gene annotations were then processed by iteratively identifying gene calls in each file, sorting them alphabetically, and enumerating the number of appearances of each gene name and displaying those counts. At the same time, a comprehensive list of all gene calls for all assemblies was tabulated and sorted while only retaining unique appearances of each gene. The list of all unique genes identified in the previous step was then used to query each list of assembly annotations using the grep function. In this case, the grep function exactly matched gene names between files and pulled the number of counts of that gene observed in each assembly or wrote a ‘0’ to indicate gene absence to a temporary file that was progressively built over time. Ultimately, a gene catalog containing a list of all unique genes and counts for those genes for each assembly was retained.

Functional gene inventories between early and mature biofilms were statistically compared for differential abundance using the DESeq2 algorithm (91). The DESeq2 algorithm (https://bioconductor.org/packages/release/bioc/html/DESeq2.html) is traditionally used for RNA-seq read count analyses but can also be used study differential abundance of genes based on a negative binomial distribution followed by p-value adjustment using the false discovery rate/Benjamini-Hochberg correction. DESeq2 internally normalizes count data by calculating the geometric mean for each gene in every sample and dividing by that mean. This normalization also corrects for library size and gene composition bias.

Metagenomic reads were taxonomically identified using MIDAS (92) for strain-level differentiation all communities. The MIDAS pipeline quantifies bacterial species abundance and strain-level genomic variation using single-nucleotide polymorphisms from shotgun metagenomes by leveraging a database of more than 31,000 prokaryotic genomes. However, the MIDAS database is geared towards studying the human microbiome, and consequently does not contain a representative database of marine microorganisms. Therefore, Kaiju (v1.8, 93) was used to complement MIDAS taxonomic annotations. Kaiju assigns each sequence in a read library to a taxon in the NCBI database with protein-level sequence homology against selectable sequence repositories. Here, we used the parameters (-x -m 9 -a greedy -e 5 -s 70 -z 8 -v) on a random subsample of reads from each read library to make a BLAST search against the non-redundant database containing bacterial and eukaryotic reads.

Metagenome assembled genomes were binned from early colonizer metagenomes using Metabat2 (94), with inclusion of only contigs greater than or equal to 2500 base pairs in length. Each MAG was processed and functionally annotated like each metagenomic assembly as described above. The quality of MAGs based on completion and contamination were assessed using CheckM (95), with a minimum percent completion of 70% and contamination less than 2%. In addition to functional metabolic annotations, taxonomic affiliation of each MAG was determined using a combination of approaches, which included taxonomic estimation with anvi-run-scg-taxonomy, anvi-run-hmms, CheckM, and GTDB-Tk (Dataset S4; 96).

### Statistical Analyses

Functional and taxonomic matrices generated from aquarium seawater cardboard, plastic, and wood communities were converted into a Bray-Curtis dissimilarity matrix and clustered using non-metric multidimensional scaling. These relative abundance data were statistically compared using a Permutational Analysis of Variance (PERMANOVA), with post-hoc multiple comparisons where applicable. Analyses of similarity (ANOSIM) were used to compare the statistical resemblance between and within levels of the factors: colonization phase, polymer type, and substrate. The relative abundance of COG categories for aquarium seawater communities and epiplastic communities were statistically compared using ANOVA or Kruskal-Wallis tests, with post-hoc multiple comparisons. The DESeq2 algorithm was used to statistically compare the differential abundance of KOfams between early colonization and mature epiplastic biofilm communities.

Carbonyl indices were statistically compared between early colonization and mature epiplastic biofilms with Mann-Whitney-Wilcoxon tests. All statistical analyses were conducted in the freely available software R (97), with packages EnhancedVolcano, ggplot2, ggpubr, pheatmap, and vegan.

## Acknowledgments

We thank Captain Charlie Howard and the crews of the *Sorcerer II* for excellent ship time services. We thank Nick Dragone for assistance with DNA extractions, Louis Kerr at the Marine Biological Laboratory (United States of America) for help with SEM imaging, and Jan van Ooijen in the nutrient laboratory at the NIOZ Royal Netherlands Institute for Sea Research.

## Data Availability

Raw sequencing data generated and analyzed in the present study have been deposited at the NCBI Sequence Read Archive under BioProject ID: PRJNA777294 and BioSample accession numbers: SAMN24661204 - SAMN24661247. Code used to create gene catalogs can be found at https://github.com/Echiostoma/. Supplemental datasets and tables can be found at https://figshare.com/s/8ced17008de131aca8d7.

## Supplementary Information for

### Supplementary Information Text

### Detailed Methods

#### Metagenome assembled genomes

The freely available software and data platform KBase (KBase, http://kbase.us/) was leveraged to bin our metagenomic data. Metagenome assembled genomes (MAGs) were mapped, binned, and inspected for quality from early colonizer read libraries using BBMap (5), Metabat2 (6), CheckM (7), and GTDB-Tk (8), respectively. Reads were mapped to metagenomic assemblies using BBMap using a minimum alignment score ratio of 0.65 and parameters (tipsearch=20, maxindel=80, minhits=2, bwr=0.18, bw=40, minratio=0.65, midpad=150, minscaf=50, quickmatch=t, rescuemismatches=15, rescuedist=800, maxsites=3, maxsites2=100, fastareadlen=500, -Xmx30g) to create a sequence alignment map (.SAM) file. The SAM file was viewed and sorted using SAMtools (samtools view -F | samtools sort - -o .BAM) and was utilized to create a sorted binary alignment map (.BAM) file for subsequent binning using Metabat2. Metabat2 was run using the syntax (--minContig 2500, --maxEdges 500). Bins established with Metabat2 were input to CheckM to inspect the quality of MAGs based on completeness and contamination. A completion percentage initially calculated by CheckM that was greater than 70% and contamination lower than 2% were required thresholds for inclusion in downstream analyses.

High-quality bins were converted into Anvi’o compatible contig databases using the anvi-gen-contigs-database command within the freely available analysis and visualization platform Anvi’o (version 7 dev). Open reading frames in contig databases were predicted using Prodigal (10) and subsequently annotated using the databases COG20 Categories and Functions (anvi-run-ncbi-cogs; 11), KOfam (anvi-run-kegg-kofams; 12), and Pfam (anvi-run-pfams; 13). These annotations were then exported (anvi-export-functions) and filtered (awk -F “\t” ‘{ if(($5 != 0) && ($5 <= .0001)) { print } }’ to include annotations with an e-value less than or equal to 0.0001. Filtered gene inventories were created using an iterative count function that outputs the final number of hits for unique gene name with the command cut -f4 $f | sort | uniq -c, where $f is the pasted file name. Taxonomic affiliation of each MAG was determined using a combination of approaches, which included taxonomic estimation with anvi-run-scg-taxonomy, anvi-run-hmms, CheckM, and GTDB-Tk (Dataset S4).

#### Synteny alignments

Amino acid sequences were exported from Anvi’o and subsequently aligned using the local pair parameterization in MAFFT (14). Aligned sequences were imported to MEGAX (15) for visualization purposes. COG20 category and function, KOfam, and Pfam annotations were programmatically matched with open reading frames and further visualized using the freely available gggenes package in R.

**Fig. S1.**
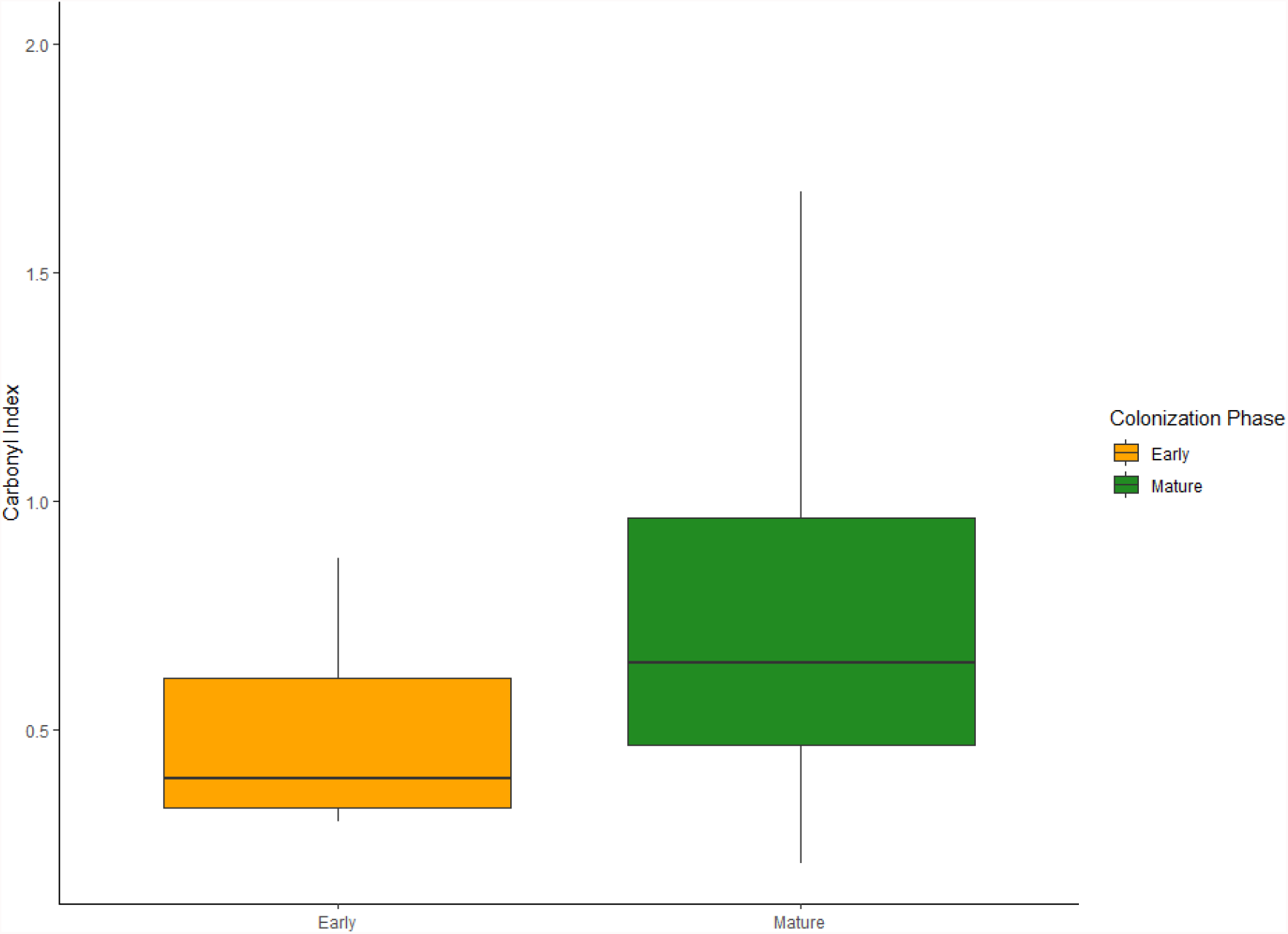
Boxplot containing carbonyl indices of plastic particles incubated during the drifter experiment (early colonization) and collected via Manta trawl (mature epiplastic biofilm). Boxes represent the interquartile range and solid horizontal black line is the median.

**Fig. S2.**
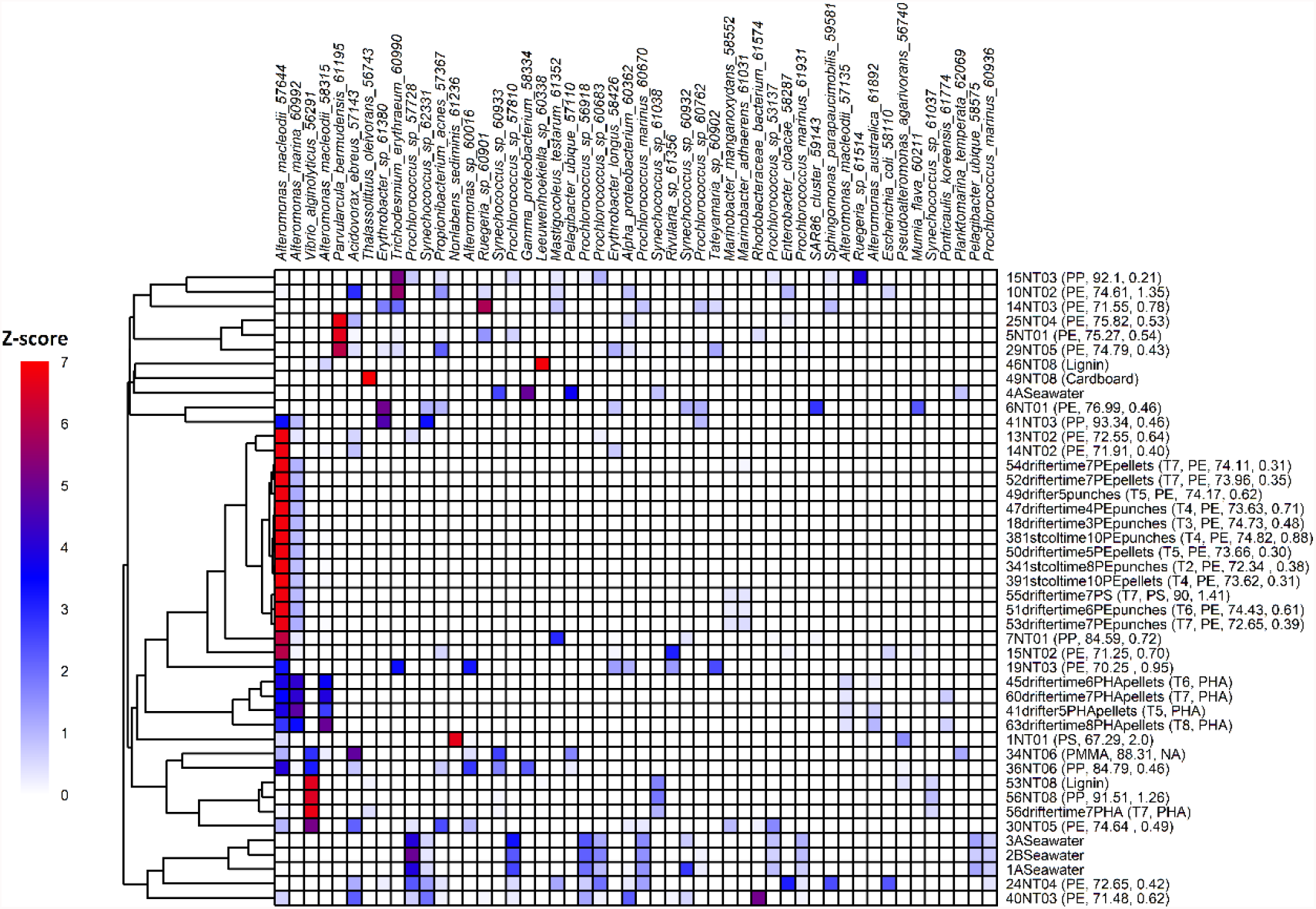
Hierarchical clustering using the top 50 bacteria species relative abundances predicted by Metagenomic Intra-species Diversity Analysis System (MIDAS) observed on cardboard, plastic, and wood samples. Z-score values for the top 50 bacteria species range between 0-7. Read library names are displayed on the right portion of this figure followed by polymer type, percent match similarity with reference spectra, and carbonyl index. PE, PMMA, PP, and PS stand for polyethylene, poly(methyl methacrylate), polypropylene, and polystyrene, respectively.

**Fig. S3.**
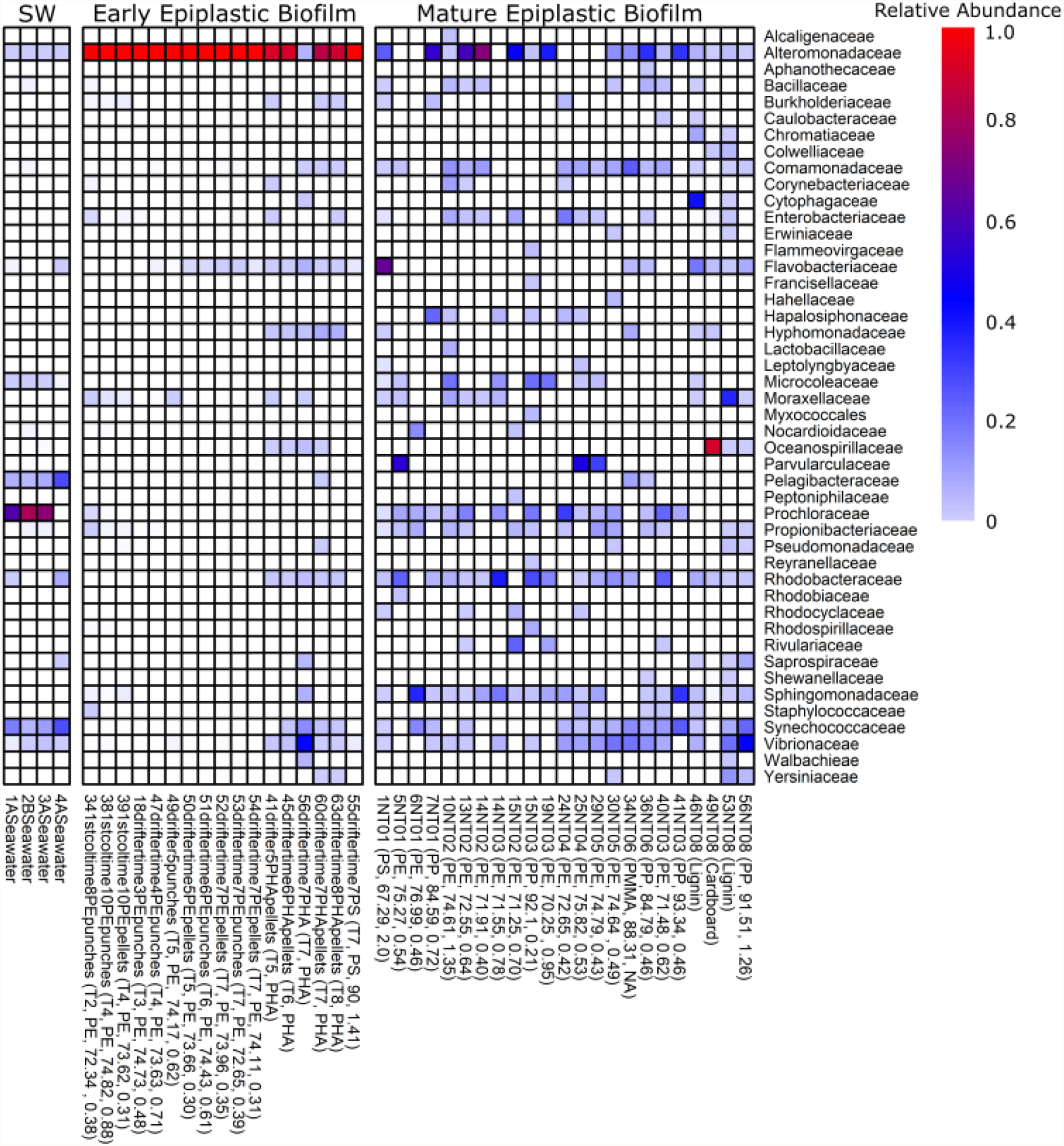
Relative abundances of prokaryote families, estimated using MIDAS, observed in seawater (SW), early colonization, and mature biofilm samples, including cardboard, plastic, and wood. Relative abundances range between 0-1. Read library names are displayed on the bottom portion of this figure followed by polymer type, percent match similarity with reference spectra, and carbonyl index. PE, PMMA, PP, and PS stand for polyethylene, poly(methyl methacrylate), polypropylene, and polystyrene, respectively.

**Fig. S4.**
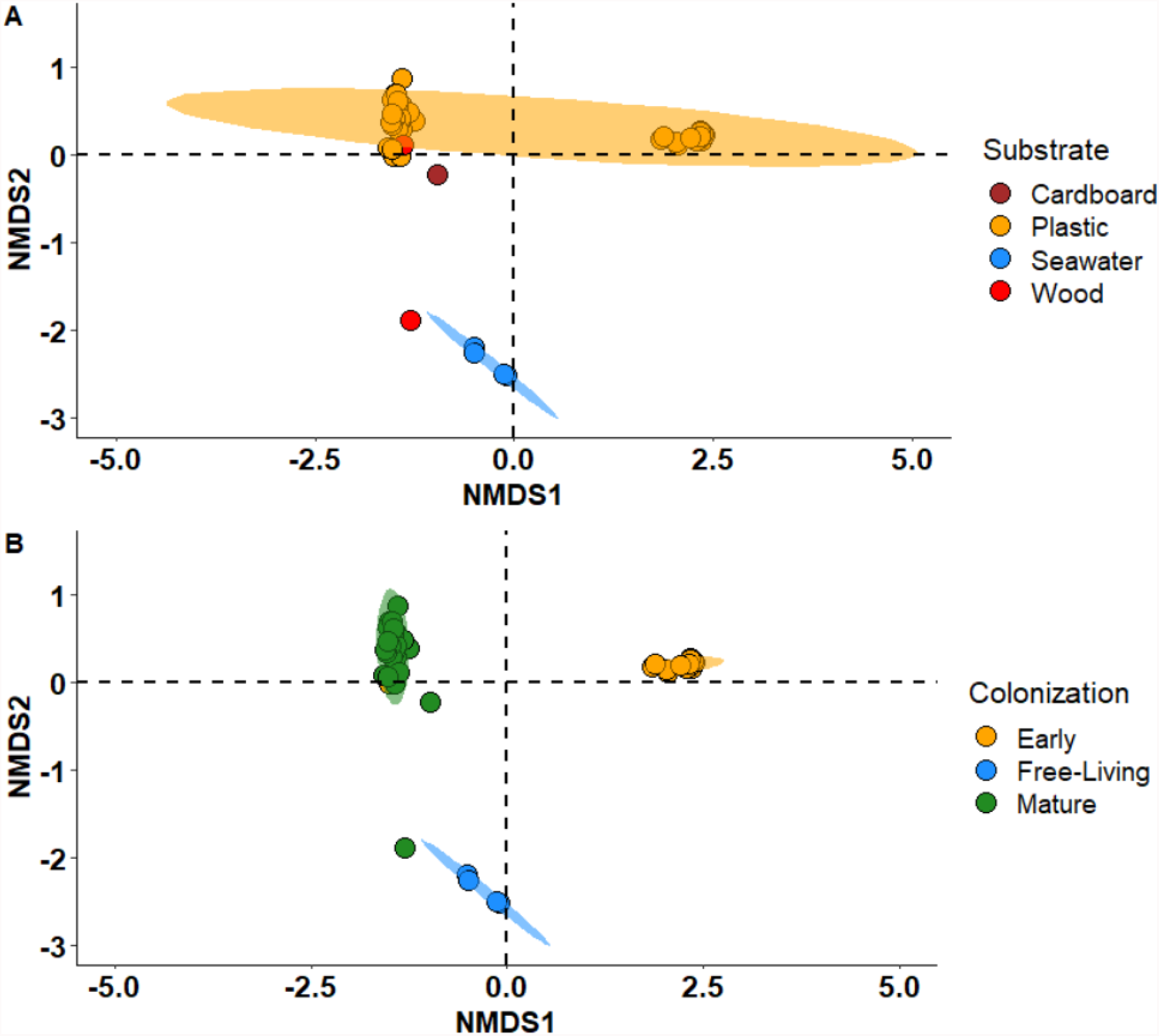
Hierarchical clustering using non-metric multi-dimensional scaling with a Bray-Curtis dissimilarity matrix containing the taxonomic resemblance by substrate (A) and colonization phase (B). Orange circle that clustered closely with mature epiplastic biofilms in panel B is PHA day 7 incubation in March 2017.

**Fig. S5.**
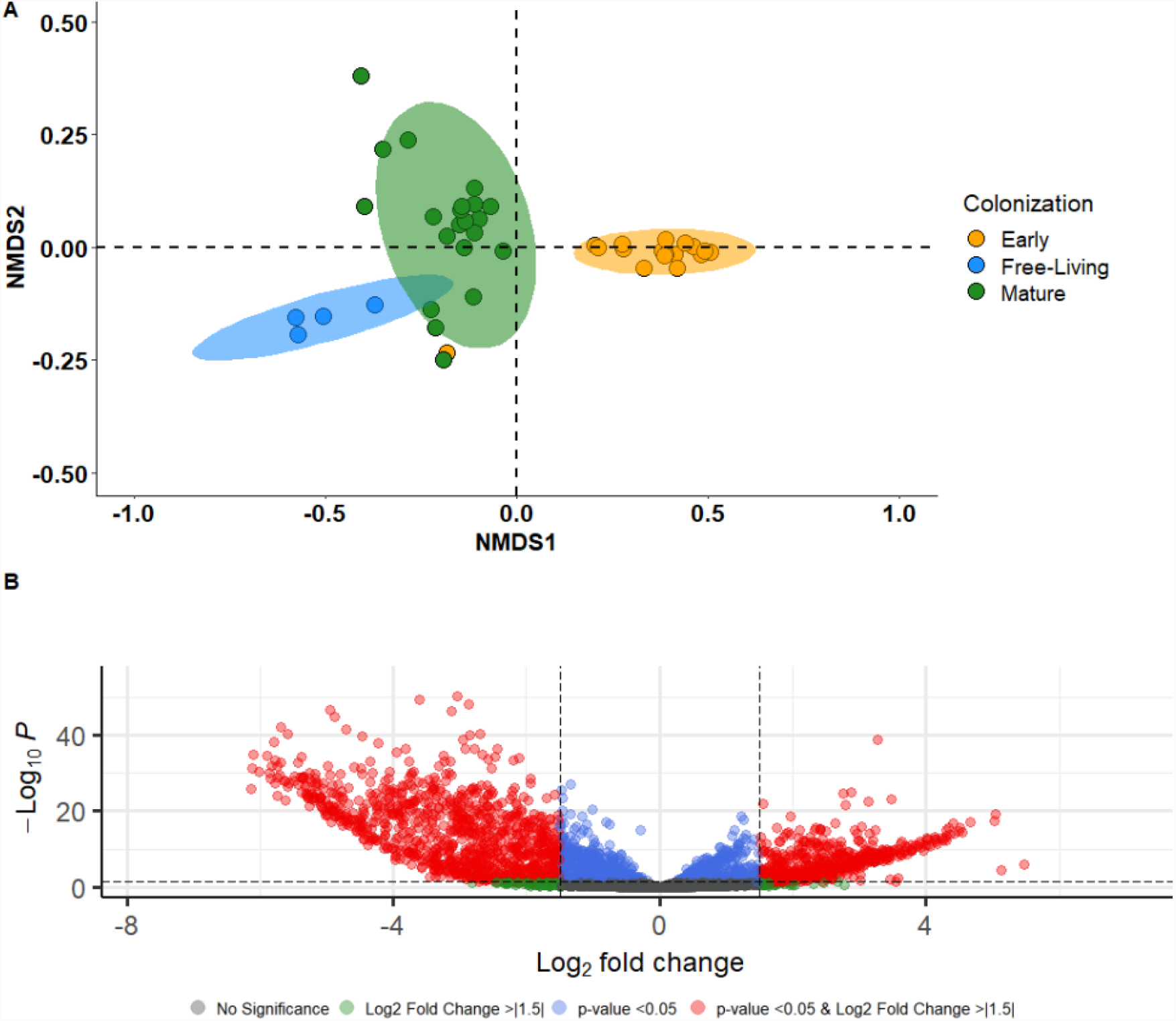
(A) Hierarchical clustering using non-metric multi-dimensional scaling with a Bray-Curtis dissimilarity matrix containing the functional metabolic resemblance (KOfam) by colonization phase. Orange circle that clustered closely with mature epiplastic biofilms in panel A is PHA day 7 incubation in March 2017. (B) Volcano plot comparing KOfam annotated genes enriched in early colonization (Log_2_ fold change <0) and mature epiplastic biofilms (Log_2_ fold changes >0).

**Fig. S6.**
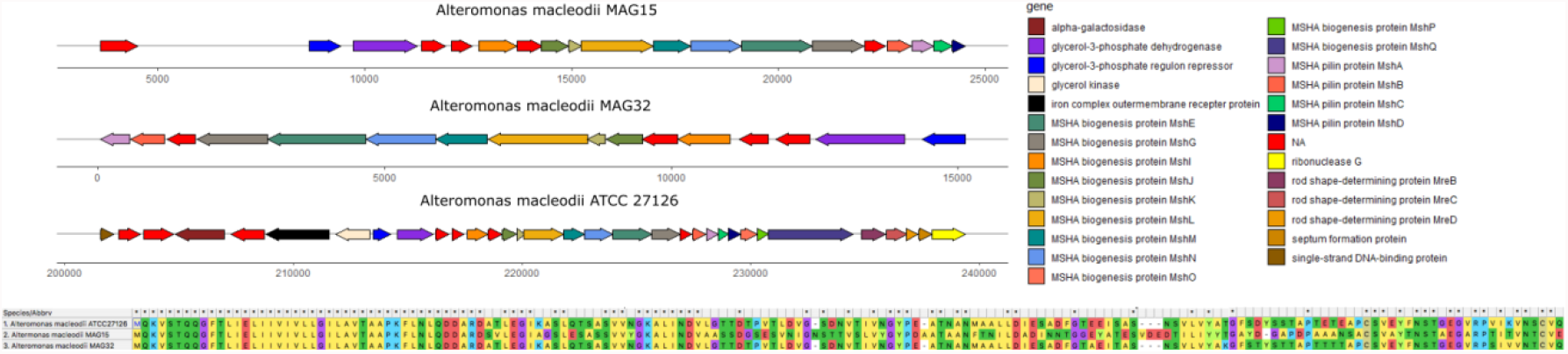
Synteny alignments of the MSHA operon and neighboring genes observed in *Alteromonas macleodii* type strain ATCC 27126 and MAGs binned from plastic biofilm metagenomes. Bottom panel is the alignment of *mshA* across genomes.

**Fig. S7.**
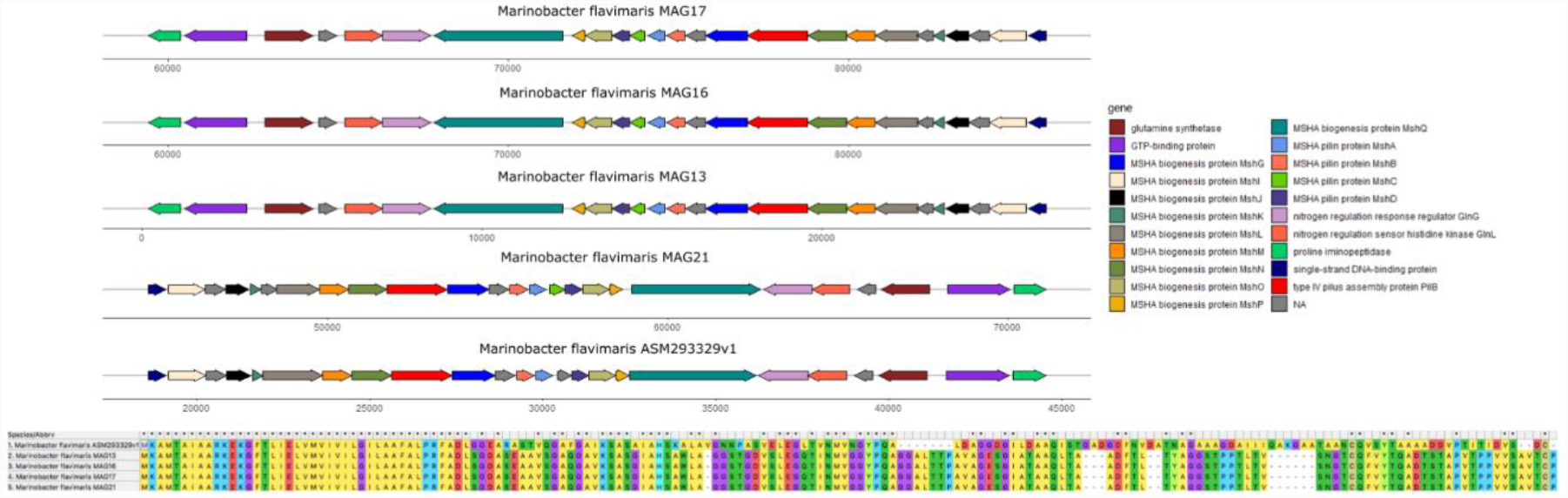
Synteny alignments of the MSHA operon and neighboring genes observed in *Marinobacter flavimaris* type strain ASM293329v1 and MAGs binned from plastic biofilm metagenomes. Bottom panel is the alignment of *mshA* across genomes.

**Fig. S8.**
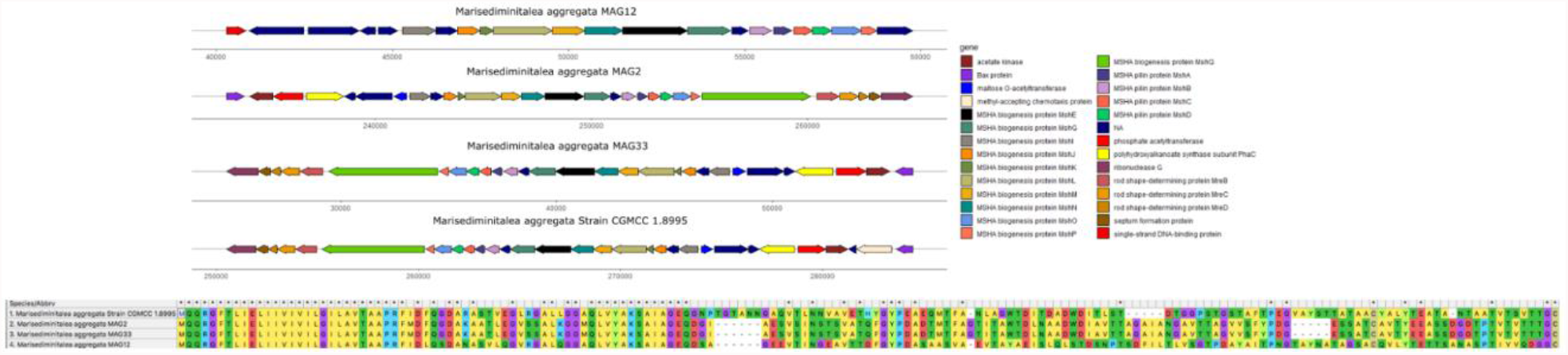
Synteny alignments of the MSHA operon and neighboring genes observed in *Marisediminitalea aggregata* type strain CGMCC 1.8995 and MAGs binned from plastic biofilm metagenomes. Bottom panel is the alignment of *mshA* across genomes.

**Fig. S9.**
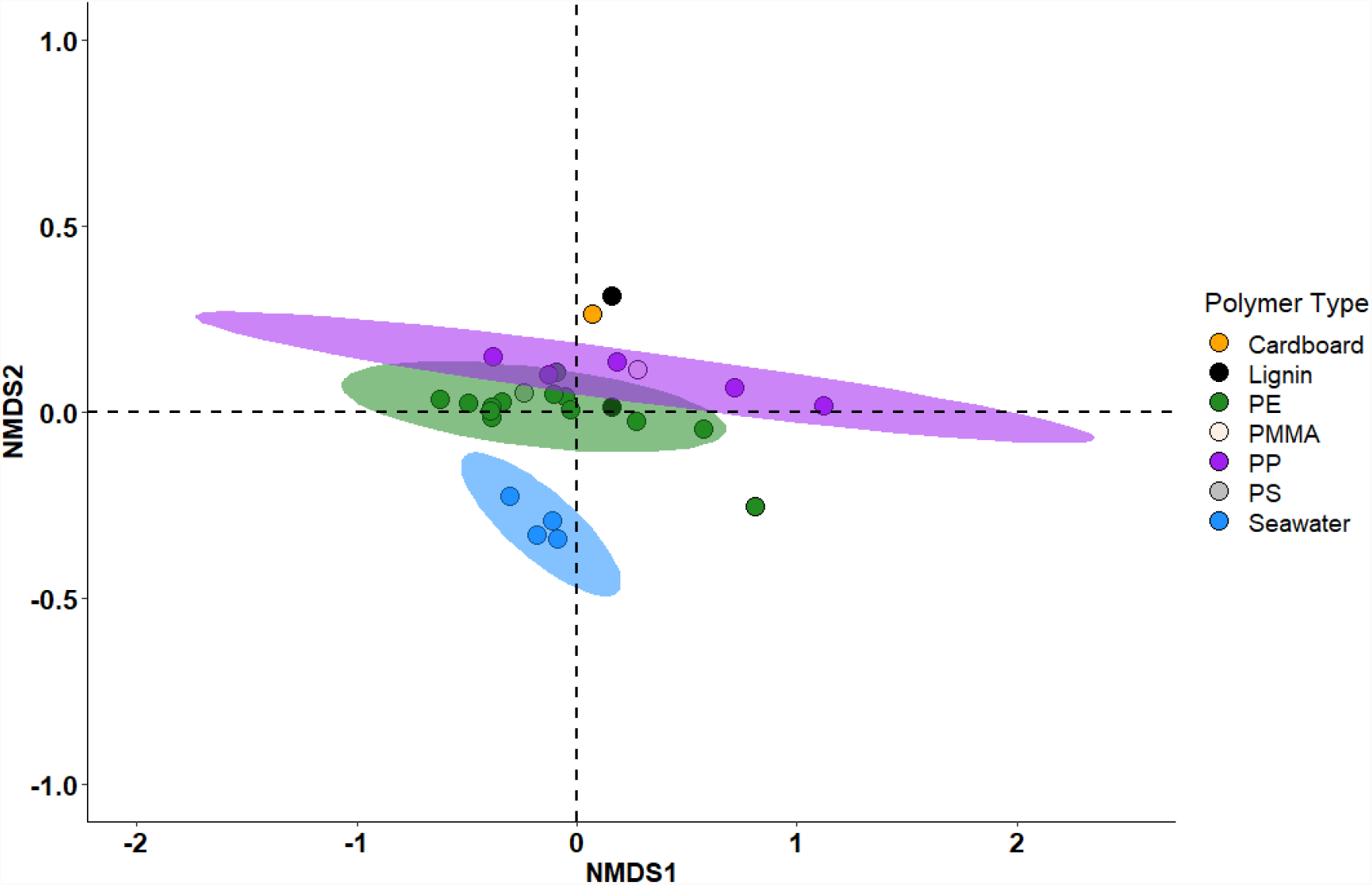
Hierarchical clustering using non-metric multi-dimensional scaling with a Bray-Curtis dissimilarity matrix containing the functional metabolic resemblance (KOfam) by polymer type/substrate of solely mature biofilm communities. PE, PMMA, PP, and PS stand for polyethylene, poly(methyl methacrylate), polypropylene, and polystyrene, respectively.

### Supplemental Table Captions

**Table S1**. Amplicon-based and whole-genome-shotgun sequencing studies of the plastisphere. WGS indicates whole-genome-shotgun sequencing.

**Table S2**. Metagenome assembly statistics.

**Table S3**. Minimum information about metagenome sequences (MIMS).

### Additional Datasets

**Dataset S4**. Annotated genomic features and taxonomic affiliation of MAGs recovered from plastics incubated during the Drifter Experiment. Dataset_S4_2022_MAG_Features

**Dataset S5**. KOfam-annotated gene inventories (counts) created for early and mature epiplastic biofilms, as well as seawater communities. Dataset_S5_2022_KOfam_Gene_Inventories

**Dataset S6**. DESeq2 algorithm outputs for KOfam annotated genes. Dataset_S6_2022_DESeq2_Outputs

**Dataset S7**. Analysis of gene copy numbers for flagella and pili involved in adhesion, chemotaxis, and motility that were observed in early colonizer MAGs. See Dataset S4 for more information on functional and taxonomic annotations of each MAG. MAG #23 was isolated from PHA day 7, which functionally and taxonomically resembled a mature epiplastic biofilm. Dataset_S7_2022_Genome_Characteristics

